# Succinct Colored de Bruijn Graphs

**DOI:** 10.1101/040071

**Authors:** Keith Belk, Christina Boucher, Alexander Bowe, Travis Gagie, Paul Morley, Martin D. Muggli, Noelle R. Noyes, Simon J. Puglisi, Rober Raymond

## Abstract

Iqbal et al. (Nature Genetics, 2012) introduced the *colored de Bruijn graph*, a variant of the classic de Bruijn graph, which is aimed at “detecting and genotyping simple and complex genetic variants in an individual or population”. Because they are intended to be applied to massive population level data, it is essential that the graphs be represented efficiently. Unfortunately, current succinct de Bruijn graph representations are not directly applicable to the colored de Bruijn graph, which require additional information to be succinctly encoded as well as support for non-standard traversal operations. Our data structure dramatically reduces the amount of memory required to store and use the colored de Bruijn graph, with some penalty to runtime, allowing it to be applied in much larger and more ambitious sequence projects than was previously possible.

## 1 Introduction

In the 20 years since it was introduced to bioinformatics by Idury and Waterman [14], the *de Bruijn graph* has become a mainstay of modern genomics, essential to genome assembly [6, 26, 21]. The near ubiquity of de Bruijn graphs has led to a number of succinct representations, which aim to implement the graph in small space, while still supporting fast navigation operations. Formally, a de Bruijn graph constructed for a set of strings (e.g., sequence reads) has a distinct vertex *v* for every unique (*k* − 1)-mer (substring of length *k* − 1) present in the strings, and a directed edge (*u,v*) for every observed *k*-mer in the strings with (*k*−1)-mer prefix *u* and (*k*−1)-mer suffix *v*. A contig corresponds to a non-branching path through this graph. See Compeau et al. [6] for a more thorough explanation of de Bruijn graphs and their use in assembly.

In 2012, Iqbal et al. [15] introduced the *colored de Bruijn graph*, a variant of the classical structure, which is aimed at “detecting and genotyping simple and complex genetic variants in an individual or population.” The edge structure of the colored de Bruijn graph is the same as the classic structure, but now to each vertex ((*k*−1)-mer) and edge (*k*-mer) is associated a list of colors corresponding to the samples in which the vertex or edge label exists. More specifically, given a set of *n* samples, there exists a set *𝒞* of *n* colors *c*_1_, *c*_2_, .., *c*_*n*_ where *c_i_* corresponds to sample *i* and all *k*-mers and (*k* − 1)-mers that are contained in sample *i* are colored with *c*_*i*_. A bubble in this graph corresponds to a directed cycle, and is shown to be indicative of biological variation by Iqbal et al. [15]. Cortex, Iqbal et al.’s [15] implementation, uses the colored de Bruijn graph to develop a method of assembling multiple genomes simultaneously, without losing track of the individuals from which (*k* − 1)-mers (and *k*-mers) originated. This assembly is derived from either multiple reference genomes, multiple samples, or a combination of both.

Variant information of an individual or population can be deduced from structure present in the colored de Bruijn graph and the colors of each *k*-mer. As implied by Iqbal et al. [15], the ultimate intended use of colored de Bruijn graphs is to apply it to massive, population-level sequence data that is now abundant due to next generation sequencing technology (NGS) and multiplexing. These technologies have enabled production of sequence data for large populations, which has led to ambitious sequencing initiatives that aim to study genetic variation for agriculturally and bio-medically important species. These initiatives include the *Genome* 10*K* project that aims to sequence the genomes of 10,000 vertebrate species [11], the *iK*5 project [10], the 150 Tomato Genome ReSequencing project [3, 16], and the 1001 Arabidopsis project, a worldwide initiative to sequence cultivars of *Arabidopsis* [31]. Given the large number of individuals and sequence data involved in these projects, it is imperative that the colored de Bruijn graph can be stored and traversed in a space-and time-efficient manner.

### Our Contribution

We develop an efficient data structure for storage and use of the colored de Bruijn graph. Compared to Cortex, Iqbal et al.’s [15] implementation, our new data structure dramatically reduces the amount of memory required to store and use the colored de Bruijn graph, with some penalty to runtime. We demonstrate this reduction in memory through a comprehensive set of experiments across the following three datasets: (1) six *Escherichia coli (E. coli)* reference genomes, (2) a set of 54 antimicrobial resistance (AMR) genes and a simulated metagenomics sample containing seven of these 54 AMR genes, and four AMR genes not contained in this set, and, (3) four plant genomes. We show our method, which we refer to as Vari (Finnish for color), has better peak memory usage on all these datasets. This observation is highlighted on our largest dataset (e.g. the plant reference genomes) where Cortex required 101 GB and Vari required 19 GB. Vari is a novel generalization of the succinct data structure for classical de Bruijn graphs due to Bowe et al. [1], which is based on the Burrows-Wheeler transform of the sequence reads, and thus, has independent theoretical importance.

In addition to demonstrating the memory and runtime of Vari, we validate its output using the *E.coli* reference genomes and AMR dataset. In particular, our experiment on the AMR dataset validates Vari’s ability to correctly identify AMR genes from a metagenomics sample, which is of paramount importance since—when expressed in bacteria—AMR genes render the bacteria resistant to antibiotics and pose serious risk to public health. Our experiments and results focus on beta-lactamases, which are genes that confer resistance to a class of antibiotics that are considered to be the last resort for infections from multi-drug-resistant bacteria [20, 25]. Our experiments demonstrate that all beta-lactamases were correctly identified and only two of the remaining 47 genes were identified to be in the sample, which had 97% and 95% sequence similarity to one of the beta-lactamases in the sample.

### Related Work

As noted above, maintenance and navigation of the de Bruijn graph is a space and time bottleneck in genome assembly. Space-efficient representations of de Bruijn graphs have thus been heavily researched in recent years. One of the first approaches was introduced by Simpson et al. [28] as part of the development of the ABySS assembler. Their method stores the graph as a distributed hash table and thus requires 336 GB to store the graph corresponding to a set of reads from a human genome (HapMap: NA18507).

In 2011, Conway and Bromage [7] reduced space requirements by using a sparse bitvector (by Okanohara and Sadakane [22]) to represent the *k*-mers (the edges), and used rank and select operations (to be described shortly) to traverse it. As a result, their representation took 32 GB for the same data set. Minia, by Chikhi and Rizk [5], uses a Bloom filter to store edges. They traverse the graph by generating all possible outgoing edges at each node and testing their membership in the Bloom filter. Using this approach, the graph was reduced to 5.7 GB on the same dataset. Contemporaneously, Bowe, Onodera, Sadakane and Shibuya [1] developed a different succinct data structure based on the Burrows-Wheeler transform [2] that requires 2.5 GB. The data structure of Bowe et al. [1] is combined with ideas from IDBA-UD [23] in a metagenomics assembler called MEGAHIT [17]. In practice MEGAHIT requires more memory than competing methods but produces significantly better assemblies. Chikhi et al. [4] implemented the de Bruijn graph using an FM-index and *minimizers*. Their method uses 1.5 GB on the same NA18507 data. In 2015, Holley et. al. [13] released the Bloom Filter Trie, which is another succinct data structure for the colored de Bruiin graph; however, we were unable to compare our method against it since it only supports the building and loading of a colored de Bruijn graph and does not contain operations to support our experiments. Lastly, SplitMEM [18] is a related algorithm to create a colored de Bruijn graph from a set of suffix trees representing the other genomes.

### Roadmap

In the next section, we describe our succinct colored de Bruijn graph data structure, generalizing Bowe et al.’s stucture for classic de Bruijn graphs [1]. Section 3 then elucidates the practical performance of the new data structure, comparing it to Cortex. Section 4 offers directions for future work.

## 2 Methods

Our data structure for colored de Bruijn graphs is based on a succinct representation of individual de Bruijn graphs that was introduced by Bowe et al. [1] and which we refer to as the BOSS representation from the authors’ initials. The BOSS representation was in turn based on an adaptation of Ferragina and Manzini’s [9] FM-indexes. Before getting to our description of the succinct colored de Bruijn graph data structure, we first describe FM-indexes and then explain the BOSS representation. Our explanation of BOSS is particularly simple and may be of independent interest to those wanting to better understand that data structure.

### 2.1 FM-indexes

Consider a string *S*. Let *F* be the list of *S*’s characters sorted lexicographically by the suffixes starting at those characters, and let *L* be the list of *S*’s characters sorted lexicographically by the suffixes starting immediately after those characters. (The names *F* and *L* are standard for these lists.) If *S*[*i*] is in position *p* in *F* then *S*[*i* − 1] is in position *p* in *L*. Moreover, if *S*[*i*] = *S*[*j*] then *S*[*i*] and *S*[*j*] have the same relative order in both lists; otherwise, their relative order in *F* is the same as their lexicographic order. This means that if *S*[*i*] is in position *p* in *L* then, assuming arrays are indexed from 0 and *≺* denotes lexicographic precedence, in *F* it is in position

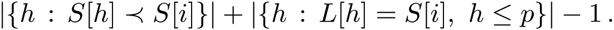

Finally, notice that the last character in *S* always appears first in *L*. It follows that we can recover *S* from *L*, which is the famous Burrows-Wheeler Transform (BWT) [2] of *S*.

The BWT was introduced as an aid to data compression: it moves characters followed by similar contexts together and thus makes many strings encountered in practice locally homogeneous and easily compressible. Ferragina and Manzini [9] realized it could also be used for indexing because, if we know the range `BWT`(*S*)[*i..j*] occupied by characters immediately preceding occurrences of a pattern *P* in *S*, then we can compute the range `BWT`(*S*)[*i′..j′*] occupied by characters immediately preceding occurrences of *cP* in *S*, for any character *c*, since

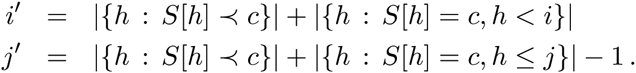

Notice *j′ − i′* + 1 is the number of occurrences of *cP* in *S*. The essential components of an FM-index for *S* are, first, an array storing |{*h*: *S*[*h*] ≺ *c*}| for each character c and, second, a rank data structure for `BWT`(*S*) that quickly tells us how often any given character occurs up to any given position^1^. To be able to locate the occurrences of patterns in *S* (in addition to just counting them), we can use a sampled suffix array of *S* and a bitvector indicating the positions in `BWT`(*S*) of the characters preceding the sampled suffixes.

### 2.2 BOSS Representation

Now consider the de Bruijn graph *G = (V, E)* for a set of *k*-mers, with each *k*-mer *a*_0_ … *a*_*k*−1_ representing a directed edge from the node labelled *a*_0_ … *a*_*k*−2_ to the node labelled *a*_1_… *a*_*k*−1_, with the edge itself labelled *a*_*k*−1_. Define the nodes’ co-lexicographic order to be the lexicographic order of their reversed labels. Let *F* be the list of *G*’s edges sorted co-lexicographically by their ending nodes, with ties broken co-lexicographically by their starting nodes (or, equivalently, by their *k*-mers’ first characters). Let *L* be the list of *G*’s edges sorted co-lexicographically by their starting nodes, with ties broken co-lexicographically by their ending nodes (or, equivalently, by their own labels). If two edges *e* and *e*’ have the same label, then they have the same relative order in both lists; otherwise, their relative order in *F* is the same as their labels’ lexicographic order. This means that if *e* is in position *p* in *L*, then in *F* it is in position

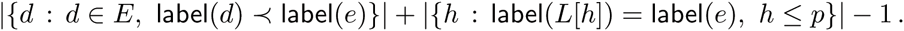

Defining the edge-BWT (EBWT) of *G* to be the sequence of edge labels sorted according to the edges’ order in *L*, so label(*L*[*h*]) = `EBWT`(*G*)[*h*] for all *h*, it follows that if we have, first, an array storing |{*d: d ∊ E*, label(*d*) *≺* c}| for each character *c* and, second, a fast rank data structure on `EBWT`(*G*) then, given an edge’s position in *L*, we can quickly compute its position in *F*.

Let *B*_*F*_ be the bitvector with a 1 marking the position in *F* of the last incoming edge of each node, and let *B*_*L*_ be the bitvector with a 1 marking the position in *L* of the last outgoing edge of each node. Given a character c and the co-lexicographic rank of a node *v*, we can use *B*_*L*_ to find the interval in *L* containing v’s outgoing edges, then we can search in `EBWT`(*G*) to find the position of the one *e* labelled *c*. We can then find e’s position in *F*, as described above. Finally, we can use *B*_*F*_ to find the co-lexicographic rank of e’s ending node. With the appropriate implementations of the data structures, we obtain the following result:

**Theorem** 1 (Bowe, Onodera, Sadakane and Shibuya, 2012). *We can store G in* (1+*o*(1))|*E*|(*lg σ* + 2) *bits such that, given a character c and the co-lexicographic rank of a node v, in 𝒪*(log log σ) *time we can find the node reached from v by following the directed edge labelled c, if such an edge exists*.

If we know the range *L*[*i*..*j*] of *k*-mers whose starting nodes end with a pattern *P* of length less than (*k −* 1), then we can compute the range *F*[*i′..j′*] of *k*-mers whose ending nodes end with *Pc*, for any character *c*
, since

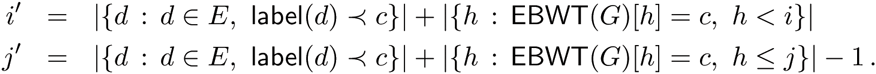

It follows that, given a node *v*’s label, we can find the interval in*L* containing *v*’s outgoing edges in 𝒪(*k* log log σ) time, provided there is a directed path to *v* (not necessarily simple) of length at least *k* − 1. In general there is no way, however, to use `EBWT`(*G*), *B_F_* and *B_L_* alone to recover the labels of nodes with no incoming edges.

To prevent information being lost and to be able to support searching for any node given its label, Bowe et al. add extra nodes and edges to the graph, such that there is a directed path of length at least *k* − 1 to each original node. Each new node’s label is a (*k* − 1)-mer that is prefixed by one or more copies of a special symbol $ not in the alphabet and lexicographically strictly less than all others. Notice that, when new nodes are added, the node labelled $^*k*−1^ is always first in co-lexicographic order and has no incoming edges. Bowe et al. also attach an extra outgoing edge labelled $, that leads nowhere, to each node with no original outgoing edge. The edge-BWT and bitvectors for this augmented graph are, together, the BOSS representation of *G*.

### 2.3 Adding Color

Given a multiset *𝒢* = {*G*_1_, …,*G*_*t*_} of individual de Bruijn graphs, we set *G* to be the union of those individual graphs and build the BOSS representation for G. We also build and store a two-dimensional binary array *C* in which *C*[*i,j*] indicates whether the ith edge in *G* is present in the jth individual de Bruijn graph (i.e., whether that edge has the jth color). (Recall from the description above that we consider the edges in *G* to be sorted lexicographically by the reversed labels of their starting nodes, with ties broken lexicographically by their own single-character labels.) We keep *C* compressed, but in such a way that we can still access individual bits quickly. If the individual graphs are similar, then most edges will appear in most graphs, so it is more natural to use 0s to indicate that edges are present and 1s to indicate that they are absent. With these data structures, we can navigate efficiently in any of the individual graphs.

Figure 1 shows an example of how we represent a colored de Bruijn graph consisting of two individual de Bruijn graphs. Suppose we are at node `ACG` in the graph, which is the co-lexicographically eighth node. Since the eighth 1 in *B*_*L*_ is *B*_*L*_[10] and it is preceded by two 0s, we see that `ACG`’s outgoing edges’ labels are in `EBWT`[8..10], so they are *A, C* and *T*. Suppose we want to follow the outgoing edge *e* labelled *C*. We see from *C*[9, 0..1] (i.e., the tenth column in *C*^*T*^) that *e* appears in the second individual graph but not the first one (i.e., it is blue but not red). There are four edges labelled *A* in the graph and three Cs in `EBWT`(*G*)[0..9], so *e* is *F*[6]. (Since edges labelled $ have only one end, they are not included in*L* or *F*.) From counting the 1s in *B_F_*[0..6], we see that *e* arrives at the fifth node in co-lexicographic order that has incoming edges. Since the first node, $$$, has no incoming edges, that means *e* arrives at the sixth node in co-lexicographic order, `CGC`.

**Figure 1:**
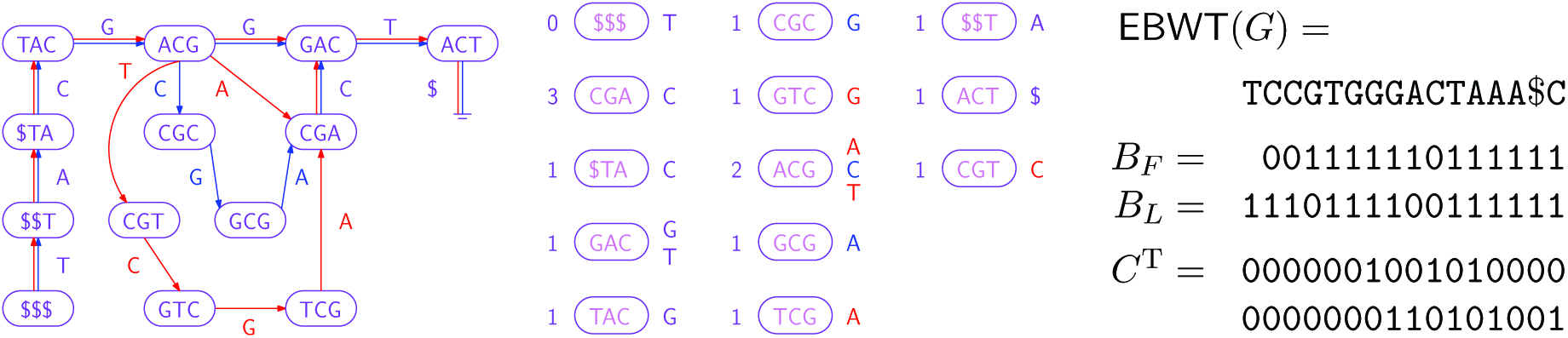
**Left:** A colored de Bruijn graph consisting of two individual graphs, whose edges are shown in red and blue. (We can consider all nodes to be present in both graphs, so they are shown in purple.) **Center**: The nodes sorted into co-lexicographic order, with each node’s number of incoming edges shown on its left and the labels of its outgoing edges shown on its right. The edge labels are shown in red or blue if the edges occur only in the respective graph, or purple if they occur in both. **Right**: Our representation of the colored de Bruijn graph: the edge-BWT and bitvectors for the BOSS representation for the union of the individual graphs, and the binary array *C* (shown transposed) whose bits indicate which edges are present in which individual graphs.

### 2.4 Implementation

We now give some details of how our data structure is implemented and constructed in practice.

#### 2.4.1 Data Structure

The arsenal of component tools available to succinct data structures designers has grown considerably in recent years, with many methods now implemented in libraries. We chose to make heavy use of the succinct data structures library (SDSL)^2^ in our implementation.

`EBWT`(*G*), the sequence of edge labels, is encoded in a wavelet tree, which allows us to perform fast rank queries, essential to all our graph navigations. The bitvectors of the wavelet tree are stored in the RRR encoding, as is the B bitvector. The rows of the color matrix, *C*, are concatenated (i.e. *C* is stored in row-major order) and this single long bit string is then stored in a RRR encoding. This reduces the size of *C* considerably because we expect rows to be very sparse (i.e. most *k*-mers are contained in most samples), and the RRR encoding is able to compress away this sparseness.

#### 2.4.2 Construction

In order to convert the input data to the format required by BOSS (that is, in correct sorted order, including dummy edges and bit vectors), we use the following process.

Our construction algorithm takes as input the set of (*k*-mer, color-set) pairs present in the input sets of reads. Here, color-set is a bit field indicating which read sets the *k*-mer occurs in^3^. We currently use the Cortex frontend to generate this set, but any *k*-mer counter capable of recording color information will suffice.

For each of these *k*-mers we generate the reverse complement (giving it the same color-set as its twin). Then, for each *k*-mer (including the reverse complements), we sort the (*k*-mer, color-set) pairs by the first *k* − 1 symbols (the source node of the edge) to give the*F* table (from here, the colors are moved around with rows of F, but ignored until the final stage). Concurrently, we also sort each *k*-mer (without the color-sets) by the last *k* − 1 symbols (the target node of the edge) to give the *L* table.

With *F* and *L* tables computed, we calculate the set difference *F - L* (comparing only the (*k* − 1)-length prefixes and suffixes respectively), which tells us which nodes require incoming dummy edges. Each such node is then shifted and prepended with $ signs to create the required incoming dummy edges (k −1 each). These incoming dummy edges are then sorted by the first *k* − 1 symbols. Let this table of sorted dummy edges be D. Note that the set difference *L - F* will give the nodes requiring outgoing dummy edges, but these do not require sorting, and so we can calculate it as is needed in the final stage.

Finally, we perform a three-way merge (by first *k* − 1 symbols) *D* with *F*, and *L - F* (calculated on the fly). For each resulting edge, we keep track of runs of equal *k* − 1 length prefixes, and *k* - 2 length suffixes of the source node, which allows us to calculate the *B*_*F*_ and *B*_*L*_ bit vectors, respectively. Next, we write the bit vectors, symbols from last column, and count of the second last column to a packed file on disk, and the colors to a separate file. The time bottleneck in the above process is clearly in sorting the *D* and *F* tables, which are of the same size, and are made up of elements of size *O(k)*. Thus, overall, construction of the data structure takes *O(k*(|*F*|log |*F*|)) time.

#### 2.4.3 Cortex’s Graph Implementation

Before discussing our experimental results, we give a brief description of the colored de Bruijn graph data structure that is implemented in the current Cortex release, which we use as a baseline to measure our performance against in the next section.

Cortex implements the colored de Bruijn graph using a hash table. Each entry in the hash table stores a (*k* − 1)-mer (vertex in the graph) as well as the following fields: the (*k* − 1)-mer labelling this vertex, coverage (an array, indexed by color), status, and edges (indicating adjacent (*k* − 1)-mers in the graph).

The (*k* − 1)-mer part of the hashtable entry is variable length, composed of multiple 64bit fields (sufficient to accommodate *k* − 1 nucleotides, represented as two bits each). The coverage information is used for error correction prior to graph construction, e.g., to remove low-coverage *k*-mers assumed to be the result of sequence errors. Later, when processing the graph, the `coverage` array is used to determine if a *k*-mer exists for a given color (coverage of 0 for a given color indicates the (*k* − 1)-mer does not exist in that sample). The `status` field is used at runtime to record whether the vertex has been previously visited, and to store other information specific to a given algorithm (e.g. bubble finding). Finally, an `edges` field is stored for each *k*-mer and each color. It is one byte in size, and each of the eight bits indicate which bases precede and follow the *k*-mer for this color. Since there are four possible predecessors and successors, one byte is sufficient.

## 3 Results

We evaluated the performance of Vari against Cortex on three different datasets, described below. Performance was evaluated on peak memory, which was measured as the maximum resident set size, and runtime, measured as the user process time. All experiments were performed on a 2 Intel(R) Xeon(R) CPU E5-2650 v2 @ 2.60 GHz server with 386 GB of RAM, and both set size and user process time were reported by the operating system. In addition to evaluating performance, we also analyzed the ability of Vari to correctly call bubbles and to accurately identify the origin of *k*-mers in a simulated metagenomics sample.

### 3.1 Datasets

Three different datasets were chosen in order to test and evaluate Vari on a variety of diverse yet realistic data types that are likely to be used as input into Vari. The first dataset contained six sub-strains of *E. coli* K-12 strain reference genomes from NCBI. Each of the genomes contained approximately 4.6 million base pairs and had a median GC content of 49.9% (1).

Our second dataset was composed of reference genomes for four different plant species: *Oryza sativa Japonica* (rice, NCBI Accession numbers: NC_008394 to NC_008405), *Solanum lycopersicum* (tomato, NCBI Accession numbers: NC_015438 to NC_015449), *Zea mays* (corn, NCBI Accession numbers: NC_024459 to NC_024468), and *Arabidopsis thaliana* (Arabidopsis, [NCBI Accession numbers: NC_003070 to NC_003076). The genome sizes and GC content were 430 Mbp and 43.42% [30], 950 Mbp and 43.42% [3, 16], 2.07 Gbp and 35.70% [27], and 135 Mbp and 47.4% [29], respectively. Hence, this represents a significantly larger dataset with more varied GC content than the *E. coli* dataset, and therefore placed more demands on both the performance and accuracy of Vari.

**Table 1:** Characteristics of the substrains of *E. coli* K-12 used to test the performance and accuracy of Vari.

| Accession Number | Sub-strain | Genome Size |
| --- | --- | --- |
| AP009048 | W3110 | 4,646,332 bp |
| CP009789 | ER3413 | 4,558,660 bp |
| CP010441 | ER3445 | 4,607,634 bp |
| CP010442 | ER3466 | 4,660,432 bp |
| CP010445 | ER3435 | 4,682,086 bp |
| U00096 | MG1655 | 4,641,652 bp |

As previously described, our third dataset contains 54 beta-lactamase genes from a custom database and a simulated metagenomics sample. We first compiled a database of known AMR genes based on sequences in the databases CARD [19], Resfinder [32] and ARG-ANNOT [12]—each of these AMR-specific databases are actively curated and contain the genetic sequences for a large variety of AMR genes. This database contains all known AMR genes, their drug resistance, and mechanism conferring resistance. We selected 54 beta-lactamase genes from this database that are known to have very high clinical and public health importance, and simulated 26,516,559 paired-end 120 bp reads from seven of the 54 beta-lactamase genes, as well as four additional AMR genes that were not included in this set of 54 genes. These latter four genes were tetracycline-resistant genes. Tetracyclines are a group of broad-spectrum antibiotics and hence, their resistance is also clinically important. This AMR dataset was used not only in the memory and time performance but also used to test the ability of Vari in identifying beta-lactamase genes from a typical metagenomic sample containing a variety of AMR genes. Table 2 contains the gene name, resistance type (beta-lactamase or tetracycline), and accession number of 11 genes that were used in simulation of the sample.

**Table 2:** List of AMR genes used to generate the simulated sample. The first seven genes were included in the the 54 beta-lactamase genes we considered for this experiment, and the remaining four were tetracycline genes. Each of the genes were approximately 1,000 bp in length and had varied GC content.

| AMR Gene | Resistance Type | Accession Number |
| --- | --- | --- |
| AmpH | beta-lactamase | AFQ67211 |
| OKP-B-4 | beta-lactamase | CAJ19612 |
| NDM-6 | beta-lactamase | AEX08599 |
| MAL-1 | beta-lactamase | CAC33434 |
| MOX-2 | beta-lactamase | CAB82578 |
| TLA-1 | beta-lactamase | ADM26831 |
| SED-1 | beta-lactamase | AAK63223 |
| TEM-1 | beta-lactamase | AFI61435 |
| TET-X | Tetracycline | AAA27471 |
| TET-X(1) | Tetracycline | ADD83116 |
| TET-C | Tetracycline | NP_387454 |
| TETR-G | Tetracycline | AAB24797 |

### 3.2 Time and Memory Usage

To compare Vari with Cortex [15], we constructed the colored de Bruijn graph, performed *bubble calling* using both data structures, and recorded the peak memory usage and runtime. Bubble calling is a simple algorithm to detect sequence variation in genomic data. It consists of iterating over a set of *k*-mers in order to find places where bubbles start and terminate. When combined with the *k*-mer color (in a colored de Bruijn graph), this enables identification of places where genomic sequences diverge from one other. A bubble is identified when a vertex has two outgoing edges. Each edge is followed in turn to navigate a path until we reach a vertex with two incoming edges. If the terminating vertex is the same for both paths, we call this a bubble. Colors for the bubbles are determined by looking at the color assignment of the corresponding (*k* − 1)-mers. Our implementation in Vari closely follows the pseudocode given by Iqbal et al. in [15]; however, it navigates the graph only in a forward direction to see if both paths converge at the same vertex, while Cortex navigates the graph backwards and forwards to find a path of adjacent vertices.

In order to test performance characteristics, this experiment was performed on all three datasets described in the previous subsection. Due to differences in the size of the datasets, the number of *k*-mers in the graph ranged from four million to over one and a half billion. As can be seen in Table 3, Vari used less than one-fifth of the peak memory that Cortex required but required greater running time. This memory and time trade-off is important in larger population level data. Given that Cortex requires 100.93 GB of space for four plant species, it would be perceptibly infeasible to run it on the i5K initiative dataset that contains the genetic sequence data for 5,000 insect species. Hence, lowering the memory usage in exchange for higher running time deservers merit in contexts where there is data from large populations.

**Table 3:**
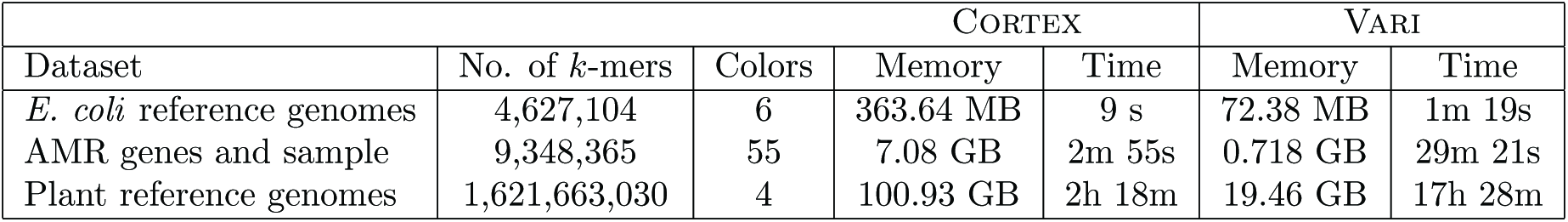
Comparison between the peak memory and time usage required to store all the *k*-mers and perform bubble finding in Cortex and Vari. *k* = 31 was used for all datasets. The peak memory is given in megabytes (MB) or gigabytes (GB). The running time is reported in seconds (s), minutes (m), and hours (h).

### 3.3 Validation on E. coli Dataset

In order to validate our data structure and test the accuracy of the bubble calling method of Vari, we compared the bubbles found by running the bubble calling algorithm on the E. coli dataset using Cortex and Vari. The bubbles outputted by each method were compared by identifying the flank preceding each bubble. Both Vari and Cortex identified 465 bubbles across all six E. coli *k*−12 substrains. This number accounts for the reverse complement bubbles found by Vari. The methods agree on 98.5% (458 / 465) of the bubbles. Thus, Vari found seven bubbles that were not identified by Cortex, which identified to be valid, and Cortex found seven bubbles not identified by Vari. These latter bubbles were missed by Vari because the addition of the reverse complement adds complexity to the graph, which changes these regions from containing a single bubble to a more complex structure. Nonetheless, our validation shows that 98.5% of the variation determined by Cortex and Vari is identical.

### 3.4 Validation on AMR Dataset

Lastly, we validated the ability of Vari to correctly identify the AMR genes contained in a metagenomics sample using a set of reference genes. Vari constructed the colored de Bruijn graph from the set of 54 beta lactamases and the simulated metagenomics sample. Hence, there were 55 unique colors in the graph because there exists one color for the metagenomic sample and one unique color for each of the 54 beta-lactamase genes. Hence, the resulting graph contains all possible *k*-mers in the dataset and the color(s) associated with each. Next, for each of the 54 genes, the unique *k*-mers were identified and the total number of these *k*-mers that were contained in the simulated sample was determined.

Table 4 in the Supplement gives the total number of each unique *k*-mers for each gene, the number of these *k*-mers that were contained in the simulated reads, and the shared *k*-mer fraction that is defined by the division of the latter two numbers. The shared *k*-mer fraction for each of the 54 genes ranged from 0.41 to 1 with a mean of 0.62. All of the seven beta-lactamase genes that were contained in the simulated sample had a shared *k*-mer fraction of 1, whereas none of the remaining 47 genes did. Of the 47 beta-lactamase genes that were not contained in the simulated sample, two had a shared *k*-mer fraction 0.98 and 0.95, however, these genes had 97% and 95% sequence similarity to one of the seven genes contained in the sample. All the remaining 45 genes had a shared *k*-mer fraction between 0.79 and 0.41. Hence, this demonstrates (on a small scale) that this use of the colored de Bruijn graph is a viable method to identify AMR genes in a metagenomics sample.

**Table 4:**
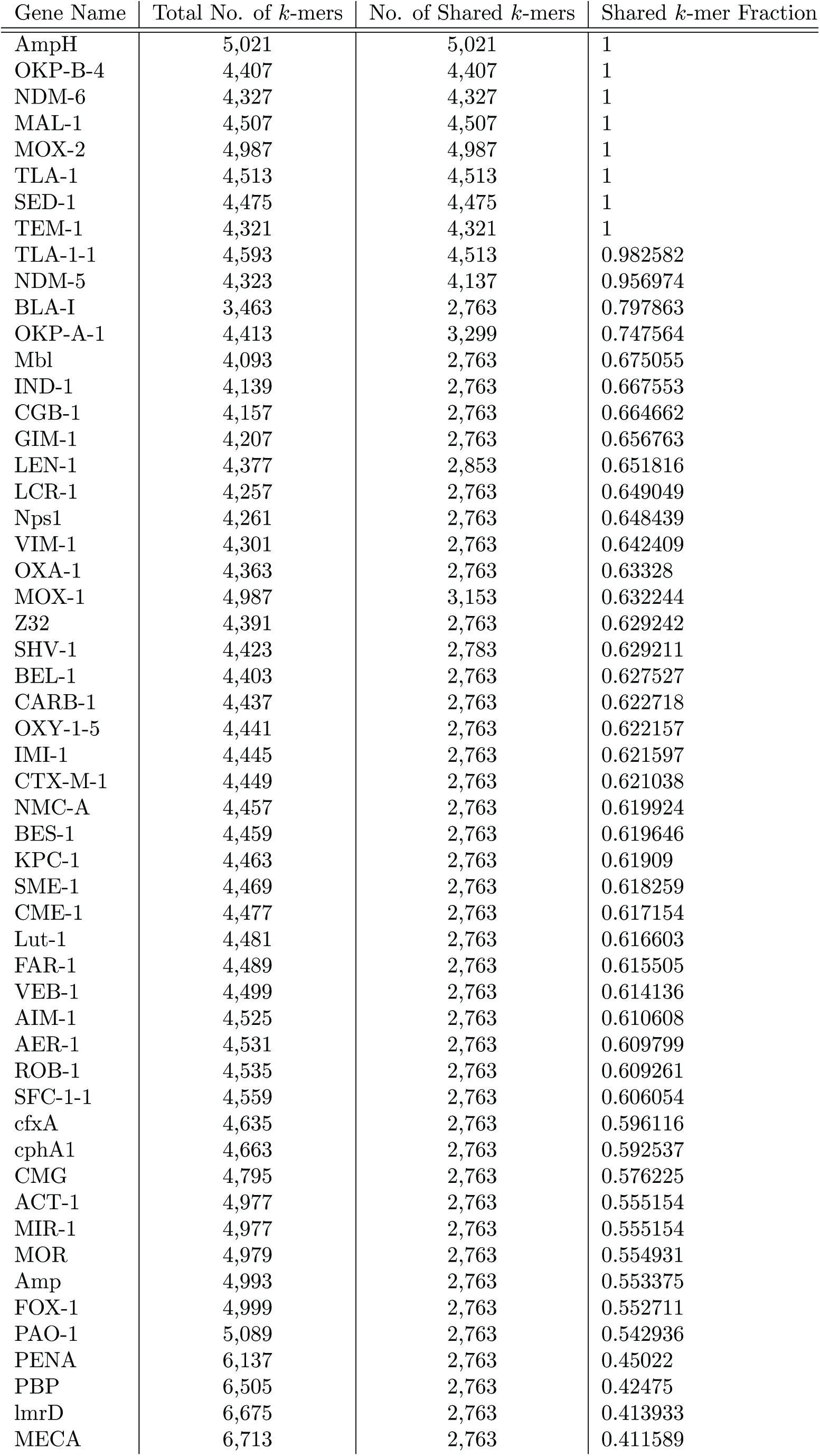
AMR gene name, number of *k*-mers in the colored de Bruijn graph, and number and proportion of *k*-mers identified in both the beta lactamase database and the simulated metagenomic sample

## 4 Concluding Remarks

We presented Vari, which is an implementation of a succinct colored de Bruijn graph that significantly reduces the amount of memory required to store and use the colored de Bruijn graph. In addition to the memory savings, we validated our approach using *E coli*. and a set of beta-lactamase genes that have a critical role in public health. Moreover, we introduced the use of colored de Bruijn graph for identifying the AMR genes within a metagenomics sample; however, as shown in our results, this use requires construction of the colored de Bruijn graph on the complete set of beta-lactamases and metagenomics sample. Nontrivial extensions to our work include (1) developing a fully scalable version of our construction algorithm that makes use of external-memory sorting, and (2) determining a succinct data structure that would allow for efficient querying of large metagenomics sample datasets. Due to the decrease in sequence costs, scientists and public health officials are increasingly moving towards a metagenomic sequence-based approach for surveillance and identification of resistant bacteria [8, 24]. A tailored colored de Bruijn graph implementation that would enable efficiently identify and comparison of AMR genes and their sequence variations across thousands of samples would be an influential method for AMR research.

## Acknowledgements

The authors would like to thank Journi Sirén from the Wellcome Trust Sanger Institute for many insightful discussions, and Zamin Iqbal for his assistance with testing Cortex.

## Supplement

Table 4 shows the results of the validation on AMR dataset as described in the Results section. The set of beta lactamase genes is listed as well as the number of *k*-mers in that genes. For each gene, we also count the number *k*-mers shared with the sample and then divide the shared *k*-mer count by the *k*-mer count to calculate the fraction. The first seven genes are in the sample so as expected their fraction is 1.

## Footnotes

1 Given a sequence (string) *S*[1,*n*] over an alphabet Σ = *{1,…, σ*}, *a* character *c ∊ Σ*, and an integer *i*, rank_*c*_(*S, i*) is the number of times that *C* appears in *S*[1,*i*].

2 https://github.com/simongog/sdsl-lite

3 In our current implementation, the color-set bitmaps were chosen to be 64 bits wide for simplicity, but can easily be extended to wider (or variable-length) bitmaps.

